# Natural occurrence and analysis of *Nosema* sp. infection in the adult population of western bean cutworm in Michigan

**DOI:** 10.1101/2022.06.29.498116

**Authors:** Dakota C. Bunn, Nicholas Miller

## Abstract

An understanding of population dynamics and insect biology is important for effective crop management strategies. Biotic factors such as pathogens play a large role on the fitness and dynamics of insect populations. Microsporidia are obligate intracellular parasites that infect more than 150 insect species and range from sublethal and chronic to fast acting and deadly. The western bean cutworm, *Striacosta albicosta* (Smith) (Lepidoptera: Noctuidae), is a pest of both corn (*Zea maize* L.) and dry beans (*Phaseolus* sp L.) that is infected by a microsporidian parasite from the genus *Nosema*. Unfortunately, little is known about the interactions between the *Nosema* sp. infecting the western bean cutworm and its prevalence and effects on the host population. This is especially true for the western bean cutworm population that has settled in the Great Lakes region over the last two decades. Using field caught samples and phase contrast microscopy, we found a 100% prevalence of infection in the adult western bean cutworm population in Michigan. No consistent trends in pathogen load were observed over the course of the western bean cutworm flight season. A weak, but statistically significant relationship was observed between male body weight and pathogen load.

## Introduction

Fitness of individuals in an insect population is influenced by multiple biotic factors such as bacteria, viruses, fungi and parasites. These factors are responsible for roughly 80% of diseases in insects and range from slow acting and non lethal to quick and extremely pathogenic (Simões et al. 2012, Solter, Becnel, and Vávra 2012, Hillyer 2016, Han et al. 2020). Microsporidia are eukaryotic, unicellular, spore forming obligate intracellular parasites found in 12 insect orders (Tokarev et al. 2020, Jordan et al. 2021). They infect more than 150 insect species and generally result in sublethal and chronic disease in infected hosts (Solter et al. 2010, Tokarev et al. 2020).

Microsporidian infection has been reported to primarily reduce feeding, fertility and life span in insect hosts as well as increase larval mortality, decrease larval size, cause delays in development, increase susceptibility to stressors and decrease flight (Dorhout et al. 2011, Solter, Becnel, and Vávra 2012, Gupta et al. 2016, Pilarska et al. 2017). Infections can be transmitted either vertically or horizontally. Vertical transmission is when the infection is transferred directly from the parent to the progeny. Whereas in horizontal transmission, individuals are infected by microsporidian spores in the environment such as contaminated water or food sources. Regardless of the method of transmission, infections with microsporidia have been both detrimental to economically important insects as well as beneficial in the control of agricultural pests (Weiss and Becnel 2014). This is especially true for the microsporidian genus *Nosema*, which are important pathogens in insects, particularly those of Lepidoptera and Hymenoptera (Solter, Becnel, and Oi 2012).

The genus *Nosema* is comprised of over 150 species and contains the majority of entomopathogenic microsporidia (Chen et al. 2009). In economically important insects such as honey bees (*Apis mellifera* (Linnaeus) (Hymenoptera: Apidae) and *Apis cerana* (Fabricius) (Hymenoptera: Apidae)), microsporidia from this genus are responsible for nosemosis, a disease that results in dysentery, shortened life spans and decreased colony size (Chen et al. 2009). The domestic silkworm, *Bombyx mori* (Linnaeus) (Lepidoptera: Bombycidae) faces a similar issue with *Nosema bombycis* (Nägeli) (Microsporidia: Nosematidae), the causative agent of pebrine disease which results in the inability of larvae to spin silkworm thread and the eventual collapse of the silkworm colony (James and Li 2012). In agriculture, *Nosema* have been proposed as a form of biological control as shown with *Nosema pyrusta* (Paillot) (Microsporidia: Nosematidae) for control of the European corn borer, *Ostrinia nubilalis* (Hubner) (Lepidoptera: Crambidae) (Lewis et al. 2009). In the lab, the sugarcane borer, *Diatraea saccharalis* (Fabr.) (Lepidoptera: Crambidae), is commonly used to produce the parasitoid, *Cotesia flavipes* (Cameron) (Hymenoptera: Braconidae) however, infection of the colony with a *Nosema* sp. pathogen does not result in the collapse of the colony. Instead it prevents the development of the parasitoid (Simões et al. 2012). Similarly, infection of the western bean cutworm, *Striacosta albicosta* (Smith) (Lepidoptera: Noctuidae), with a *Nosema* sp. pathogen has been suggested to increase the ability of the western bean cutworm to tolerate *Bacillus thuringiensis* toxins (Helms and Wedberg 1976).

The western bean cutworm is a univoltine pest of corn (*Zea mays* L.) and dry beans (*Phaseolus* sp L.) that is infected with a microsporidian parasite from the *Nosema* genus (Helms and Wedberg 1976, Su 1976, Dorhout 2007, Dyer et al. 2013). Native to the western Great Plains region of the United States, it has been expanding its range eastward since the late 1990s and is now established in at least twenty-five states and four Canadian provinces (Hoerner 1948, Hagen 1962, Jeschke 2018). The western bean cutworm is particularly a problem in corn where feeding provides entry for molds and other plant pathogens, increasing the risk of diseases and mycotoxin contamination (Peairs 2014). In corn, large infestations are able to reduce yield by 30 to 40%, and while less serious than in corn, dry beans yield is reduced by 8 to 10% (Peairs 2014, Difonzo et al. 2015). Management of this pest has proven difficult and while field research is ongoing, difficulty in rearing it in the lab has made it challenging to research the organism in depth and unfortunately, research into the effects of the *Nosema* sp. infection on the western bean cutworm is limited (Dyer et al. 2013, Smith et al. 2019).

Previous research into the interaction between the western bean cutworm and its associated *Nosema* sp. has focused on the life cycle of the microsporidian parasite and its prevalence of the infection based on location (Su 1976, Dorhout 2007). Su (1976) observed that samples collected between 1970 and 1975 in Nebraska, before the western bean cutworm range expansion, had a 100% prevalence (the number of infected individuals in a population). Dorhout studied samples ranging from Wyoming to Ohio with a focus on central Iowa during the western bean cutworm expansion in 2004 and 2006, observing less than a 57% prevalence in their Iowa samples with a varying range of intensity (the concentration of parasites infecting the host) (Dorhout 2007). However, to date, no studies have been performed on this interaction using populations from Michigan following its establishment in the Great Lakes region in 2006 (DiFonzo and Hammond 2008).

Understanding the population structure, abundance and dynamics of insect pests is important for the development of effective pest management strategies (Rusch et al. 2010, Combs et al. 2019). While unable to determine a reason, Bunn et al. (2021) found that in Michigan two population peaks were observed annually for the western bean cutworm. Infection, pathogens and disease play important roles in population dynamics with microsporidian infections in invertebrate host populations fluctuating seasonally (Araújo-Coutinho et al. 2004, Herren et al. 2020). Unfortunately, understanding of the prevalence and effects of *Nosema* sp. infection on the fitness and population dynamics of the western bean cutworm is limited (Smith et al. 2019).

In order to start to understand these effects, we hypothesized that the prevalence and intensity of the *Nosema* sp. infection in the adult population of western bean cutworm changes throughout the flight season in Michigan. To test this hypothesis, we captured adult western bean cutworm samples from central Michigan in 2017 and 2018, extracted the *Nosema* spores, and using phase contrast microscopy determined the prevalence and intensity of infection. We then performed further analysis to determine how sex, trap date, and size may interact with the intensity of infection.

## Materials and Methods

### Western Bean Cutworm Sample Collection

Adult western bean cutworms were collected in central Michigan throughout the flight season (mid-July through mid-August) in 2017 and 2018 (Bunn et al. 2021). Samples were selected from six universal pheromone moth traps (Great Lakes IPM Inc., Vestaburg, MI) that were used in 2017 and the three Quantum Black Light Traps (Leptraps LLC, Georgetown, KY) used in 2018. In 2017 traps were baited with western bean cutworm sex pheromone lures (Scentry Biologicals Inc., Billings, MT). All traps were spaced between 2.8 and 31.4 km apart to ensure the traps were operating independently and contained a piece of insecticidal strip (Hercon Vaportape) to prevent moths from escaping or becoming damaged due to prolonged activity in the enclosed space. Traps were checked every morning and moths were placed in 50 mL centrifuge tubes containing 95% ethanol, returned to the lab, and stored at -80°C until they were processed for spore extraction.

### Spore Extraction

For each trap date, spore samples were extracted from 30 randomly selected adult moths spread evenly across all traps (pheromone traps in 2017 and light traps in 2018) whenever possible. Individual abdomens of samples were dissected at the junction between the abdomen and thorax, and their weights were determined. Each abdomen was placed into a 1.5 mL microcentrifuge tube and homogenized in 1 mL of double distilled (dd) water with a tissue homogenizer (Raun et al. 1960, Solter, Becnel, and Vávra 2012). Once ground the samples were left for 72 hours at 4°C to allow time for the host cells to lyse and then centrifuged at 1500 g for 20 minutes to pellet the spores. The supernatant was removed, and the pellets were resuspended in 1 mL of dd water. Each sample was individually transferred and diluted to a total volume of 10 mL with dd water before being filtered into individual 15 mL tubes with a 10 mL luer lock syringe fitted with a 5.0 μm polypropylene syringe filter (Tisch Scientific, Cleves, Ohio). The filtered material was centrifuged for 20 minutes at 1500 g and the supernatant was removed (Han et al. 2019).

The samples were then washed to remove any additional cellular debris. To do so, they were resuspended in 1 mL of PBS-Tween20 (0.3%) and centrifuged for 20 minutes at 1500 g, after which the supernatant was removed, and the step was repeated before performing the same washing steps an additional two times with PBS 1X (Han et al. 2019). Finally, the spores were resuspended in PBS (1X). For resuspension, 0.1 mL of PBS was used for every 3 mg of the original tissue weight (Raun et al. 1960, Solter, Becnel, and Vávra 2012).

### Confirmation of Microsporidia

#### DNA Extraction

An additional adult western bean cutworm was selected at random, and the spores were extracted as previously described with the protocol only varying at the final resuspension, where the sample was resuspended in 1 mL of PBS (1X). DNA was extracted as described by Pombert et al. (Pombert et al. 2013). First, following resuspension, host DNA was removed by adding 10 μL of DNase I (10 mg/mL) (Sigma-Aldrich, St. Louis, Missouri) and 15 μL of filtered MgCl_2_ solution (1M) to the sample and incubated at room temperature for 15 minutes under rotation. Then 60 μL of EDTA (0.5 M) was added and the sample was mixed by inversion before being centrifuged at 1,500 g for 20 min. The supernatant was discarded, and the spore pellet was washed by being resuspended in 1 mL of PBS (1X) and centrifuged at 1,500 g for 20 min. This was repeated for a total of 6 washes. Finally, the washed spores were resuspended in 1 mL of PBS (1X) and stored at 4°C until later use.

Following the removal of host DNA, the spores were pelleted by centrifugation (3000 g, 15 min) and the supernatant was discarded. Subsequently, 1 μL of proteinase K (50 g/L) and 300 μL of MasterPure gram positive cell lysis solution (Lucigen, Middleton, Wisconsin) were added to the pelleted spores. The spores were resuspended by pipetting and vortexing before being transferred to a tube containing 200 μL of glass beads (150-212 m diameter). Bead beating was performed for 30 seconds at 2,500 rpm before the spore sample was incubated at 65°C for 18 minutes with additional bead beating performed every 6 minutes for 30 seconds at 2,500 rpm. The sample was cooled and incubated at 37°C for 30 minutes upon the addition of 2 μL of RNase A (5 μg/μL, Lucigen, Middleton, Wisconsin).

Once the incubation was complete, the sample was placed on ice for 5 minutes to deactivate the enzyme and for the glass beads to settle. The supernatant was then transferred to a new 1.5 mL tube. MPC protein precipitation reagent (10 μL, Lucigen, Middleton, Wisconsin) was added to the new tube and vortexed for 10 seconds before being centrifuged at 4°C for 10 minutes (≥17,000 g). The supernatant was transferred to a new 1.5 mL microcentrifuge tube before 500 μL of cold 100% isopropanol was added. The tube containing the supernatant and the isopropanol was inverted 40 times manually before once again being centrifuged at 4°C for 10 minutes (≥17,000 g). The supernatant was removed, and the remaining pellet was rinsed twice with 300 μL of cold ethanol (70%) before being allowed to air dry for 7 minutes. Following this, 35 μL of ultra-pure water was added to the pellet and the DNA was stored at 4°C overnight to resuspend. Once the DNA was resuspended, the DNA was quantified using a dsDNA High Sensitivity quantification kit (Invitrogen, Waltham, MA) and a Qubit 3.0 (Invitrogen, Waltham, MA) according to the manufacturer’s instructions.

#### PCR

General microsporidian primers targeting the small subunit ribosomal RNA and the internal transcribed spacer, ss530f (5’-GTGCCAGCMGCCGCGG-3’) and ss1047r (5’-AACGGCCATGCACCAC-3’) (Weiss and Becnel 2014), were used to perform a PCR on the extracted spore DNA along with a positive control of *Encephalitozoon intestinalis* DNA and negative control of molecular biology grade water. The PCR reactions were performed in 50 μL of 1X Green GoTaq Buffer (Promega, Madison, Wisconsin), containing 1.5 mM MgCl_2_, 1.25 units DNA polymerase (Promega, Madison, Wisconsin), 0.2 mM of each dNTP, 1 μM of each primer, and 1 ng of template DNA. The PCR conditions were an initial denaturation at 95°C for 2 minutes, followed by 40 cycles of denaturation (95°C for 30 seconds), annealing (55°C for 30 seconds), and extension (72°C for 2 minutes). Following these cycles, a final extension was performed at 72°C for 5 minutes. PCR products were visualized on a 1% agarose gel made with TAE (1X) and ethidium bromide (1 μg/mL) for 90 minutes (55V, 400 mAmp) along with a 100 bp size standard ladder (New England Biolabs, Ipswich, Massachusetts).

#### Imaging and Counting

A hemocytometer was used to image each sample under a 10X objective using phase contrast microscopy (Nikon Eclipse Ts2) and a Canon EOS REBEL T3i digital camera. The presence or absence of any spores found in a 0.20 mm^2^ area of the hemocytometer was used to determine the prevalence of the infection in the population. To determine the intensity of each infection, the number of spores in the 0.20 mm^2^ area were counted and the spores per milligram of tissue were calculated according to:

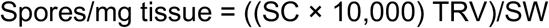

Where SC is the spore count obtained from the hemocytometer, TRV is the total resuspension volume (0.1 mL for every 3 mg tissue weight), and SW is the original abdomen sample weight in mg.

## Statistical Analysis

All statistical analysis was performed using R Statistical Software (R Core Team 2022). Spore count data was log transformed and a Shapiro-Wilk (SW) normality test was performed on both the non-transformed and the transformed data. A non-parametric Scheirer-Ray-Hare (SRH) test (Mangiafico 2016) was then performed using the transformed data for the spore count vs sex and trap day, spore count vs trap year and trap day (Males), and spore count vs sex and trap location. Finally, a linear model fitting and analysis of variance was performed for the abdomen size vs spore count in both males and females.

## Results

Primary confirmation of microsporidian spores was obtained using phase contrast microscopy, shown in Figure 1, and were observed to be ovate and consistent with the average size of 4.45 μm reported by Su in 1976. The DNA extraction resulted in 35 μL of DNA at a concentration of 16.3 ng/μL with a size greater than 10.0 kb, indicating a high molecular weight (Figure 2). Secondary confirmation of microsporidian spores was performed using PCR with the extracted DNA and universal microsporidian primers resulting in a PCR product between 450 to 500 bp (Figures 3).

**Figure 1.**
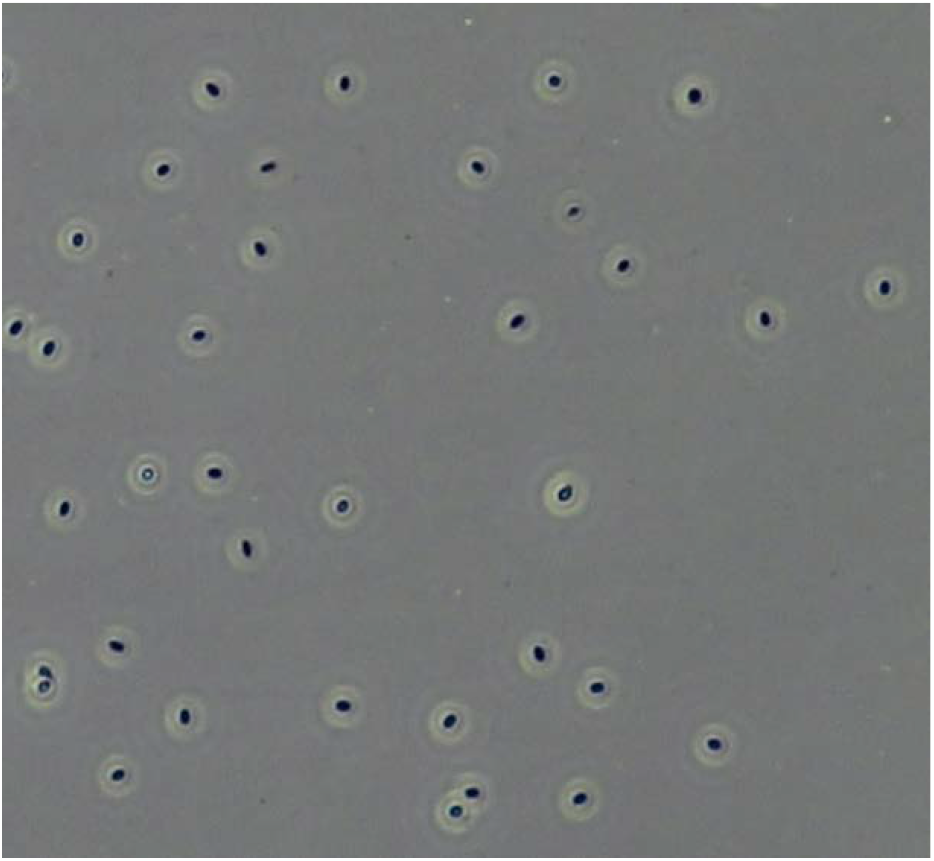
Image of *Nosema* sp. spores extracted from adult male western bean cutworm under phase contrast microscopy.

**Figure 2.**
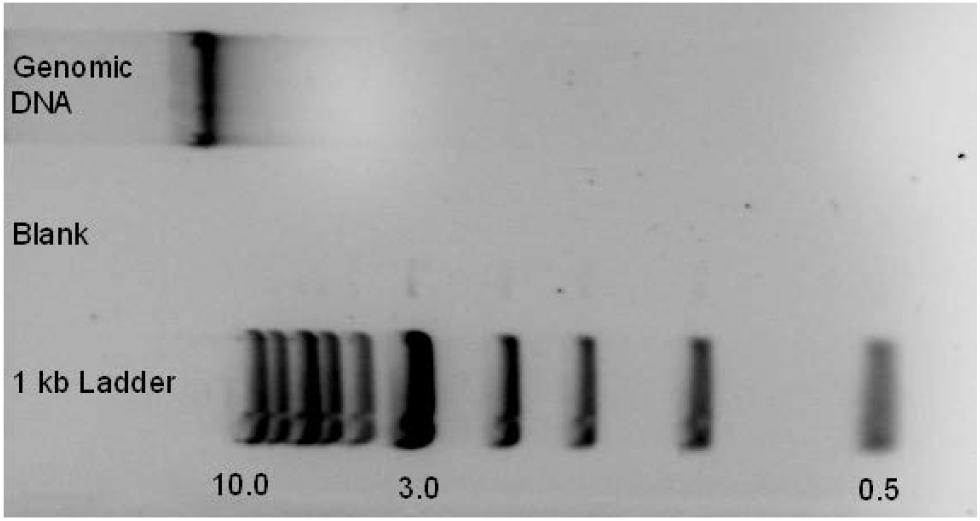
Ethidium bromide gel showing the confirmation of high molecular weight DNA extracted from *Nosema* sp. spores. Lane 1 contains a 10kb ladder, lane 2 was left blank, and lane 3 contains the *Nosema* sp. DNA.

**Figure 3.**
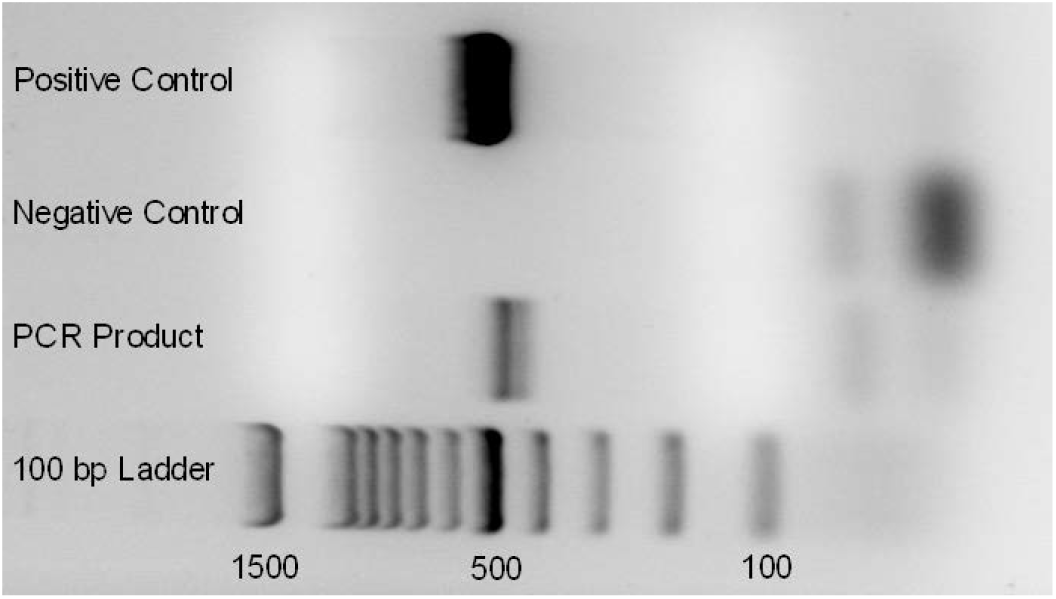
PCR results of general microsporidian primers. Lane 1 contains a 100bp ladder, lane 2 depicts the product of the primers and the extracted *Nosema* sp. DNA, lane 3 is the negative control and lane 4 is the positive control.

Of the 3,264 moths captured across the two study years (Bunn et al. 2021), 22% (738) of the moths were used to determine the occurrence of *Nosema* sp. infection in the population. One hundred percent of the observed population was found to be infected, with infection intensity ranging between 330 to 886,000 spores/mg tissue.

In 2018, 403 moths were randomly selected with 362 males and 41 females. The distribution of the raw spore count data was determined to be significantly non-normal (SW normality test: W = 0.32074, p-value < 10^−4^), and the data were log transformed to improve visualization, although it did not normalize the distribution (SW normality test: W = 0.95894, p-value < 10^−4^). Mean log spore counts ranged from 7.71 ± 1.14 to 11.64 ± 0.87 as shown in Figure 4. Female moths showed slightly higher intensity, and a significant difference in intensity based on sex was observed (SRH test: H = 7.254, df = 1, p-value = 0.007), although the difference was modest (Figures 4 and 5). Significant differences were observed in intensity based on trap day (SRH test: H = 62.036, df = 15, p-value < 10^−4^) and no interaction between trap day and sex in intensity was found (SRH test: H = 13.176, df = 9, p-value = 0.15).

**Figure 4.**
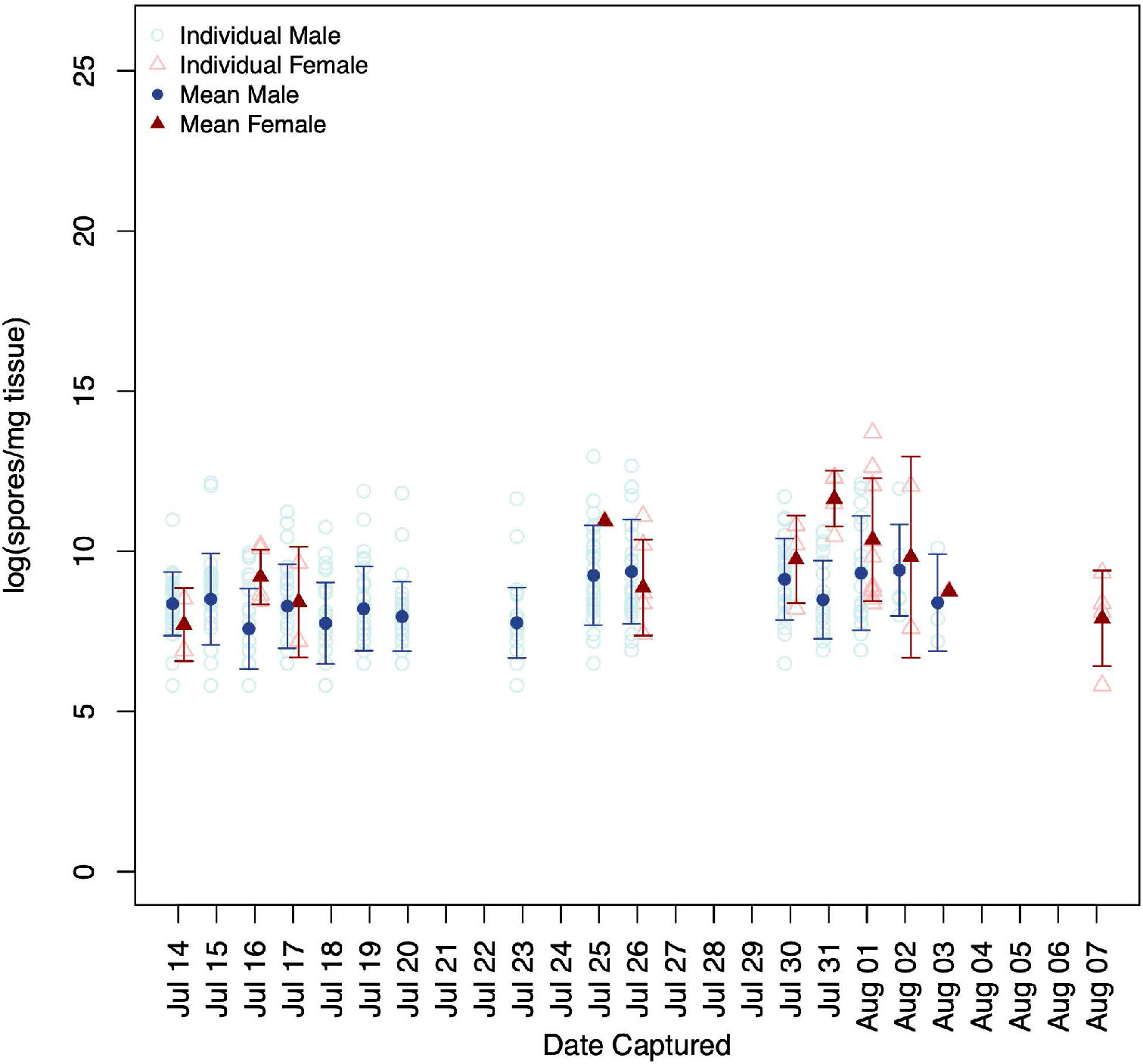
Parasite load in light trap captured adults in 2018 throughout the flight season. Males are denoted with circles while females are denoted with triangles. Mean values are shown as filled shapes with the appropriate standard deviation.

**Figure 5.**
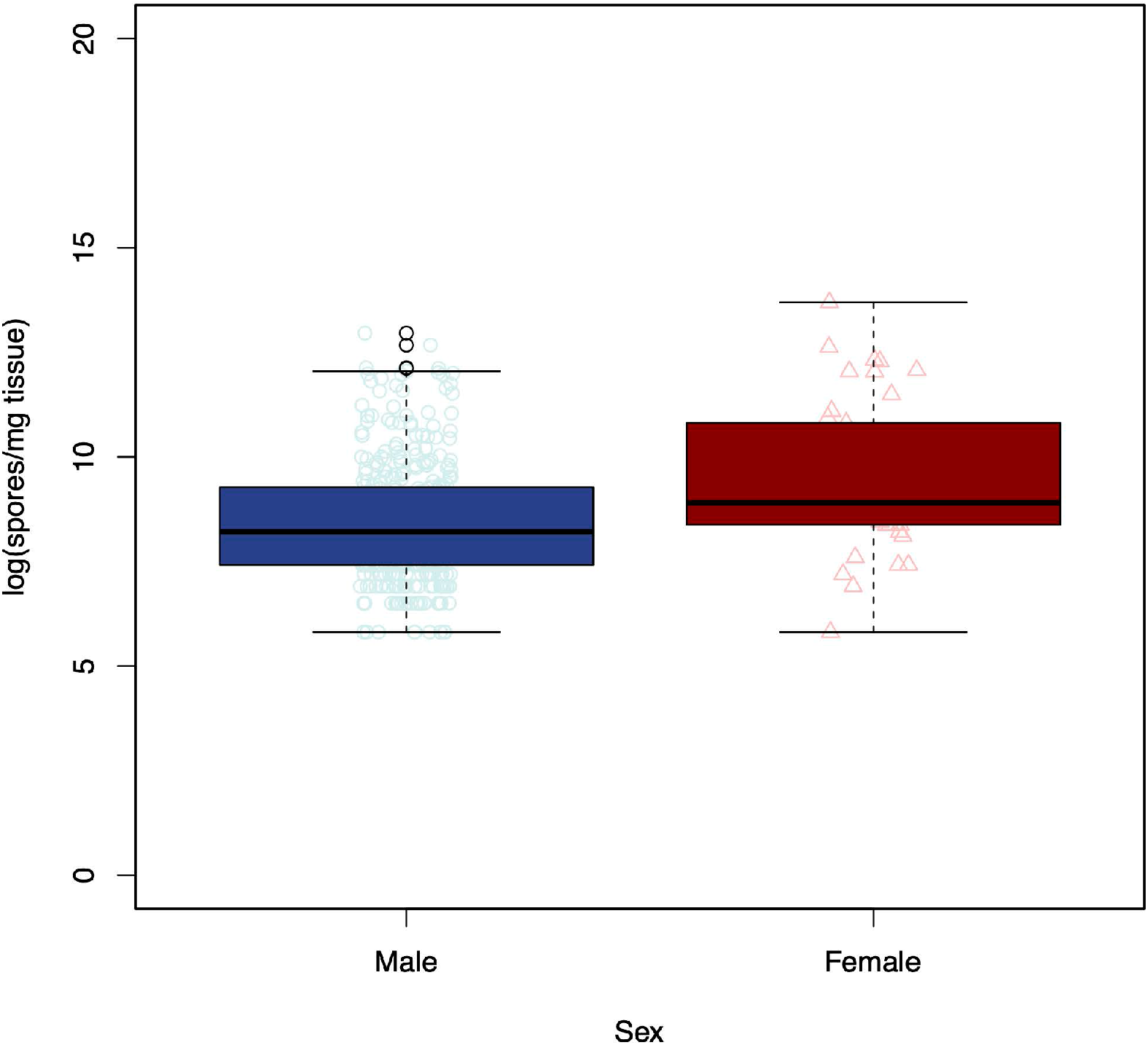
Parasite load in light trap captured males and females in 2018.

To compare trap days between the two years, 333 males were randomly selected from 2017 and analyzed against the males previously selected in 2018. This study was confined to males because in 2017, trapping was performed primarily with pheromone traps due to technical issues as noted in Bunn et al. (2021). The distribution of the raw spore count data for these samples were determined to be significantly non-normal (SW normality test: W = 0.39084, p-value < 10^−4^) and the data were log transformed for visualization purposes (SW normality test: W =0.95464, p-value < 10^−4^). Mean intensities ranged from 7.57 ± 0.52 to 10.17 ± 1.57, with significant differences observed between trap days (SRH test: H = 71.785, df = 22, p-value < 10^−4^), trap year (SRH test: H = 17.76, df = 1, p-value < 10^−4^) and the interaction between the two (SRH test: H = 17.588, df = 8, p-value < 0.02) (Figure 6).

**Figure 6.**
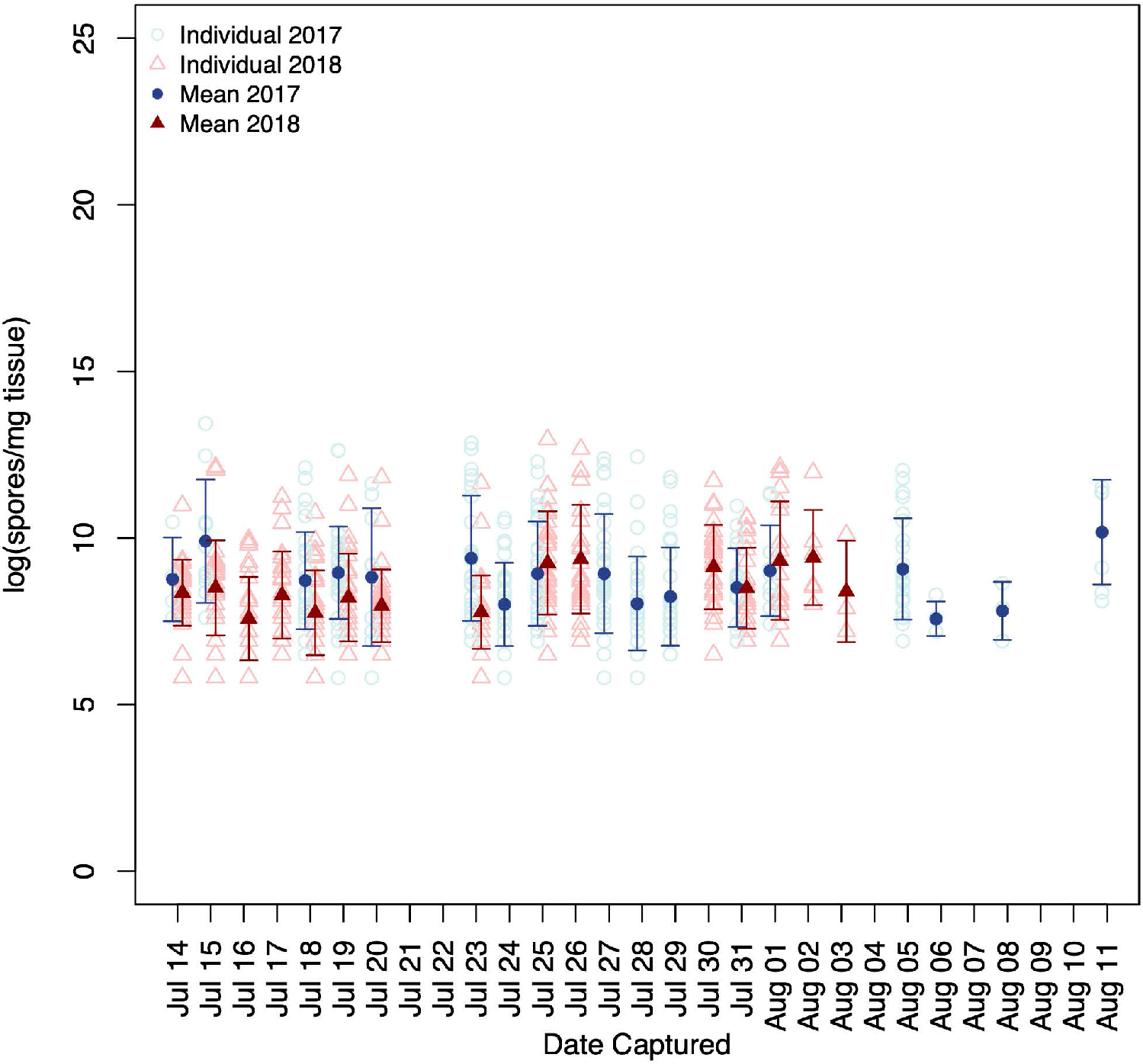
Parasite load in captured males throughout the flight seasons in 2017 and 2018. Males from 2017 are denoted with circles while males from 2018 are denoted with triangles. Mean values are shown as filled shapes with the appropriate standard deviation.

When comparing the log spore count (spores/mg of tissue) with the size of the moths abdomen, a linear model fitting resulted in a significant effect on the spores per milligram of tissue based on the original sample weight for males as shown in Figure 7 (R^2^ = 0.06, p-value = 5.16e-11).

**Figure 7.**
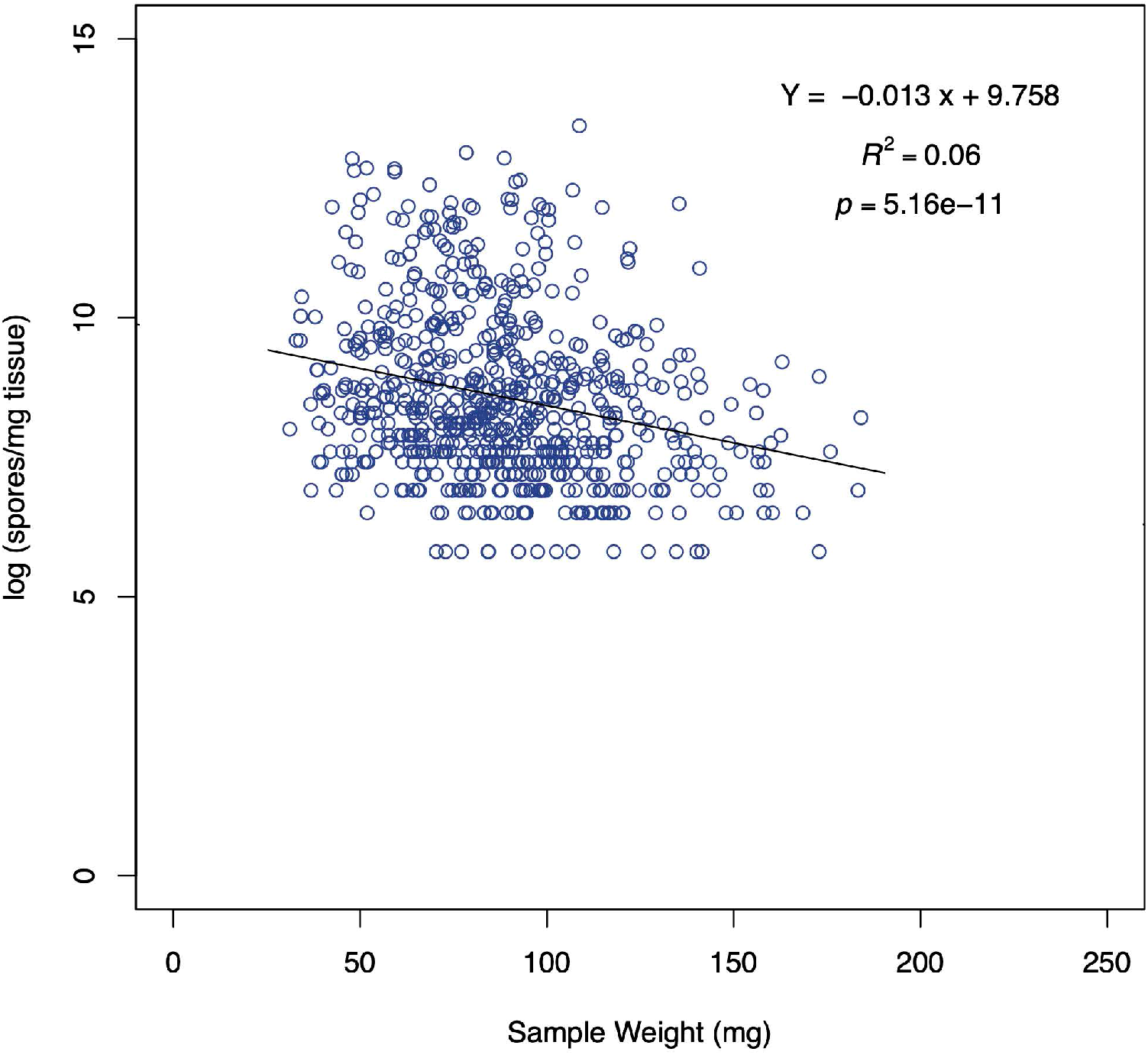
Parasite load in males vs sample weight.

Performing the same analysis with female samples, no significant effect on spores per milligram of tissue was observed, as shown in Figure 8 (R^2^ = -0.02, p-value = 0.6).

**Figure 8.**
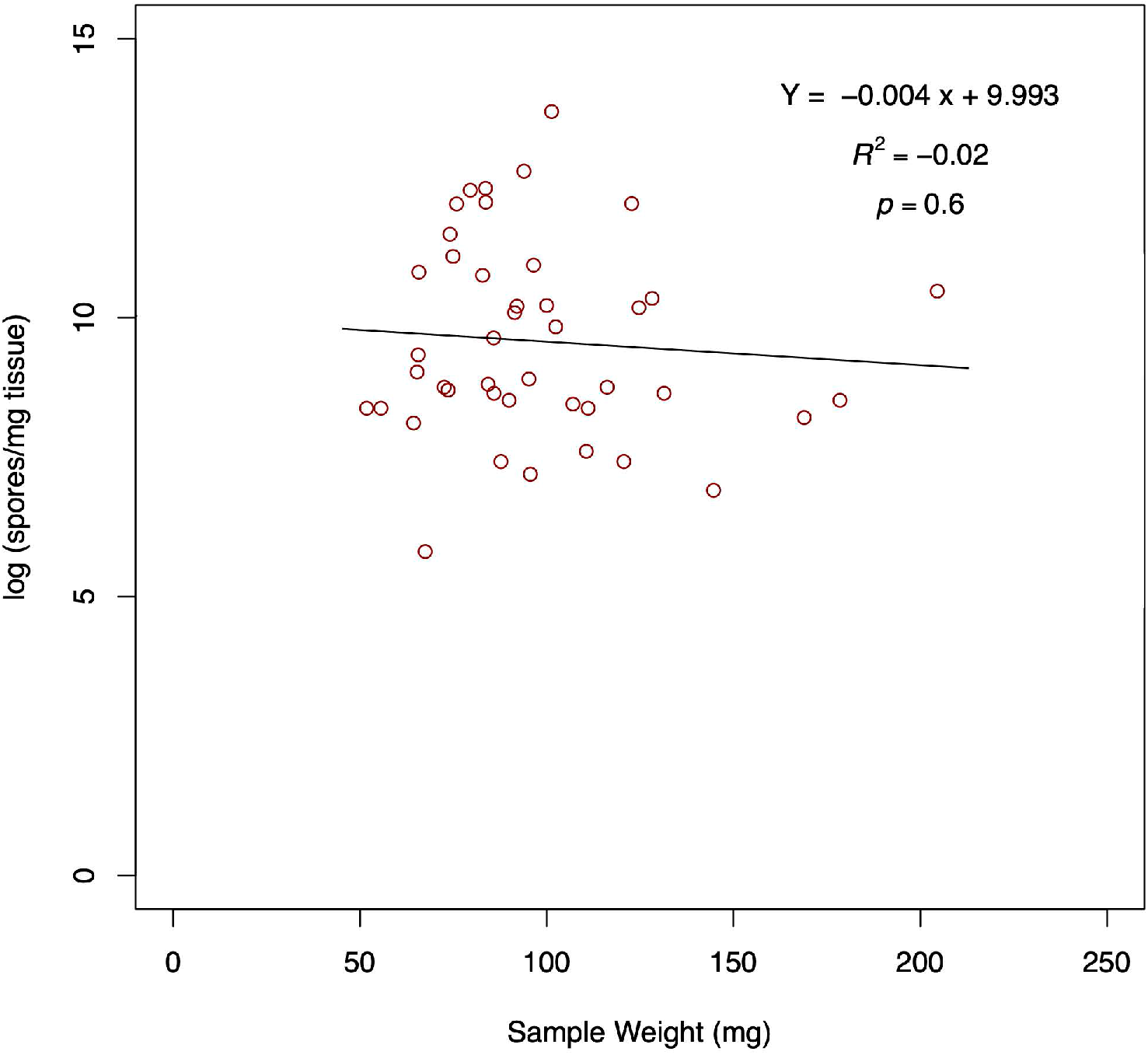
Parasite load in females vs sample weight.

## Discussion

Of the 778 tested samples, 100% were infected with microsporidian spores, with intensity ranging from 330 to 886,000 spores/mg tissue. This suggests that there is a 100% prevalence of microsporidian infection in the Michigan western bean cutworm population. These findings are similar to reports by Su in 1976 on Nebraska caught samples, where all specimens were infected with *Nosema* spores (Su 1976).

A similar study, performed on samples collected between Wyoming and Ohio in 2004 and 2006, did not show 100% prevalence of the *Nosema* sp. in the population (Dorhout 2007). The aim of Dorhout’s study was to assess prevalence across a range of locations representing both the ancestral and expanding host range of the western bean cutworm, whereas our study and the study performed by Su (1976) focused on more localized samples. While we cannot confirm the reason for the discrepancies in *Nosema* sp. prevalence, it has previously been observed that, in general, the low virulence of microsporidian pathogens allows for an increase in prevalence in the population over time (Biganski et al. 2021, LeBrun et al. 2022). A possible explanation for the difference in results may be that during the range expansion by the western bean cutworm, not all individuals were infected with the microsporidian parasite. However, as the expanded population stabilized, the prevalence increased over the past 20 years resulting in the 100% prevalence observed today.

When attempting to rear western bean cutworm in the lab, the *Nosema* sp. infecting the larvae has been suggested to potentially be one of the challenges faced as larvae are particularly susceptible to disease (Dyer et al. 2013, Smith et al. 2019). The high mortality observed in lab reared larvae is somewhat surprising considering the 100% prevalence of *Nosema* sp. infection observed in the adult population. One explanation of this may be the forced proximity of the host organisms. The western bean cutworm is not a social insect compared to other *Nosema* genus hosts such as honeybees or the forced sociability of the domestic silkworm, where infections result in high rates of mortality (Chen et al. 2009, 2013, Bjørnson and Oi 2014, Gupta et al. 2016). It is therefore unclear if the mortality observed in the western bean cutworm larvae is due to the progression of the infection itself or the close confines required in a laboratory setting which may be forcing an increase in exposure to the pathogen and severity of the disease.

In 2018, a significant difference in infection load between males and females, as well as between the trap days, was observed as shown in figures 4 and 5. While it has not been well studied in Lepidoptera, several studies have noted differences in intensity based on sex, as well as various sex dependent responses to microsporidian infection (Wilson 1978, Solter et al. 2002). This has also been observed in Hymenoptera and Coleoptera (Boohene et al. 2003, Bjørnson and Oi 2014). The observation of differences in intensity between different trap dates was expected as microsporidian infection in other species fluctuates seasonally (Abdel-Baki et al. 2009, Herren et al. 2020). No obvious pattern for intensity over time was found and while we observed interactions between the two factors of trap day and sex, no general pattern was discernable.

In males captured in 2017 and 2018, no pattern was immediately evident despite a significant difference in infection load between trap years, trap days and the interaction between the two as shown in figure 6. While the results may be statistically significant, insect pathogens, like human pathogens, are known to vary annually and seasonally (Abdel-Baki et al. 2009, Herren et al. 2020, LeBrun et al. 2022).

When comparing abdomen size to infection intensity we observed a significant effect between abdomen size and intensity in males (Figure 6). While the effect of microsporidia infection on adult size has not been well studied, various studies in Lepidoptera have found that larval size tends to decrease due to microsporidian infection (Rusch et al. 2010, Solter, Becnel, and Oi 2012, Kermani et al. 2013, Bjørnson and Oi 2014, Gupta et al. 2016) and when examining the size of adult insects, it is important to remember that large larvae tend to develop into large adults (Nijhout et al. 2006, Kivelä et al. 2020). The same effect was not observed in females. Sex based responses to microsporidian infection have been noted in other insects, including Lepidoptera, which may be the case here (Wilson 1978, Solter et al. 2002, Bjørnson and Oi 2014).

Overall, our data suggest that the *Nosema* sp. infecting the western bean cutworm behaves as do the majority of microsporidia, in that it results in sublethal and chronic disease in the host organism. While it was not examined in our study, microsporidia infections are known to remain sublethal in the population until the end of the adult life cycle, upon which the adults become susceptible to the infection due to the addition of other environmental stressors (Solter, Becnel, and Oi 2012, Bjørnson and Oi 2014, Weiss and Becnel 2014). This may also be the case for the *Nosema* sp. infected western bean cutworm. The interaction between infected western bean cutworm and transgenic corn expressing *Bt*-toxins warrants further study. Bunn et al. (2021) found that in central Michigan, where transgenic hybrids are the most common mechanism of control for the western bean cutworm in corn, the majority of adult moths developed on corn as larvae (Bunn et al. 2021). When coupled with previous studies that suggest exposure of infected western bean cutworm to *Bt*-toxins provokes a beneficial response in the host and decreases microsporidian infection (Helms and Wedberg 1976), it is apparent that more research is required into this interaction to ensure that the most efficient pest management strategies are being developed and employed.

## Acknowledgements

We would like to thank Leellen Solter, PhD for her training in microsporidia microscopy and identification. We would also like to thank Jean-Francois Pombert, PhD and Anne Caroline Mascarenhas dos Santos, MS for access to their BSL-2 facilities and assistance with experimental troubleshooting.

